# Heritable gene expression variability governs clonal heterogeneity in circadian period

**DOI:** 10.1101/731075

**Authors:** K.L. Nikhil, Sandra Korge, Kramer Achim

## Abstract

A ubiquitous feature of circadian clocks across life forms is its organization as a network of coupled cellular oscillators. Individual cellular oscillators of the network often exhibit a considerable degree of heterogeneity in their intrinsic periods. While the interaction of coupling and heterogeneity in circadian clock networks is hypothesized to influence clock’s entrainability, our knowledge of mechanisms governing network heterogeneity remains elusive. In this study, we aimed to explore the principles that underlie inter-cellular period variation in circadian clock networks (clonal period-heterogeneity). To this end, we employed a laboratory selection approach and derived a panel of 25 clonal cell populations exhibiting circadian periods ranging from 22 h to 28 h. We report that while a single parent clone can produce progeny clones with a wide distribution of circadian periods, heterogeneity is not entirely stochastically driven but has a strong heritable component. By quantifying the expression of 20 circadian clock and clock-associated genes across our panel, we found that inheritance of different expression patterns in at least three clock genes might govern clonal period-heterogeneity in circadian clock networks. Furthermore, we provide preliminary evidence suggesting that epigenetic variation might underlie such gene expression variation.

## INTRODUCTION

The majority of life forms on earth exhibit ∼24 h (circadian) behavioural and physiological rhythms generated by endogenous time-keeping mechanisms - circadian clocks. In addition to driving such endogenous rhythms, circadian clocks facilitate synchronization of organisms’ rhythms to daily and seasonal changes in the environment to enhance their survivability, thereby functioning as an adaptive mechanism (Kumar, 2017). The fundamental basis of circadian rhythm-generation across all life-forms are cell-autonomous molecular oscillations comprising evolutionarily conserved auto-regulatory transcription-translation feedback loops (TTFL) (Dunlap, 1999). In higher organisms, such cell-autonomous clocks often function as a network of coupled oscillators which in unison drive circadian rhythms (Bell-Pedersen *et al*., 2005). Welsh and co-workers first reported that neurons within the suprachiasmatic nucleus (SCN; the master pacemaker in the hypothalamus of mammals) are surprisingly heterogeneous in their intrinsic periods of circadian firing pattern (Welsh *et al*., 1995). Subsequent studies revealed that such period-heterogeneity is not restricted to the SCN alone, but is also characteristic of mammalian peripheral clock cells (Nagoshi *et al*., 2004; Leise *et al*., 2012) as well as of clock cells in Drosophila (Sabado *et al*., 2017) and plants (Yakir *et al*., 2011; Muranaka and Oyama, 2016). The ubiquity of this network feature suggests that heterogeneity may be functionally relevant for circadian clocks (Jagota *et al.,* 2000; Schaap *et al*., 2003; Gonze *et al*., 2005; Bernard *et al*., 2007; Inagaki *et al*., 2007; VanderLeest *et al*., 2007; Gu *et al*., 2016, 2019), thus likely being a substrate for natural selection. Interestingly, the observed period-heterogeneity among circadian clock cells within an organism cannot be entirely attributed to functionally different cell types as cells of the same subtype (clonal cells) also exhibit such variation (Nagoshi *et al*., 2004; Leise *et al*., 2016). Clonal-heterogeneity or clonal-phenotypic variability is common in biology and can stem from various external factors such as stochastic changes in the microenvironment or internal factors like stochastic partitioning of cellular components during cell-division or stochasticity in gene expression (Neildez-Nguyen *et al*., 2008; Brock, Chang and Huang, 2009; Huang, 2009; Altschuler and Wu, 2010; Geiler-Samerotte *et al*., 2013; Roberfroid, Vanderleyden and Steenackers, 2016; Evans *et al*., 2018). In this study, we aimed to explore the possible mechanisms underlying clonal-heterogeneity of circadian period in human circadian oscillator cells.

We hypothesised that clonal period-heterogeneity in mammalian cells is due to a) stochastic variation (Geva-Zatorsky *et al*., 2006; Chang *et al*., 2008; Brock, Chang and Huang, 2009; Frank and Rosner, 2012) and/or b) heritable variation (Dubnau and Losick, 2006; Gordon *et al*., 2009, 2013). Since the term ‘stochastic’ is used in the context of both non-heritable (external noise and gene expression noise) as well as heritable gene expression variation (epigenetic stochasticity), for the rest of this manuscript we define ‘stochasticity’ as any non-heritable variation (both internal and external). To test the two hypothesis outlined above, we employed a laboratory selection approach and derived a panel of 25 clonal cell lines (from a common ancestral/founding culture) exhibiting a range of periods between 22h and 28h. We observed that the period-heterogeneity among progeny clones stemming from a single parent cell is not entirely stochastic but has a substantial heritable component. We then measured expression of 20 clock and clock-associated genes in our panel and observed that variation in gene expression levels of at least three clock genes (transcription factors) might underlie clonal period-heterogeneity. Furthermore, we provide preliminary evidence that epigenetic variation might govern the observed clonal-variation in gene expression.

## RESULTS

### Clonal period-heterogeneity is not entirely stochastically driven but is largely inherited

Is the variation in period among individual circadian oscillator cells just due to intrinsic and/or extrinsic stochastic noise? Or is there a heritable component? To test this, we single-cell cloned a ‘founding culture’ of U-2 OS cells (an established model of peripheral circadian clocks) harboring a *BMAL1*-luciferase reporter construct (Maier *et al*., 2009). Upon reaching confluence, the period of bioluminescence rhythms from these progeny cultures was determined by live-cell bioluminescence recording. As expected, we observed a distribution of circadian periods (23.5 h - 27.5 h; Figure 1a top panel). We repeated this protocol for several ‘assay-generations’ by each time selecting short and long period cultures as ‘parents’ for the next assay-generation (Study outline in Supplementary Figure S1).

**Figure 1:**
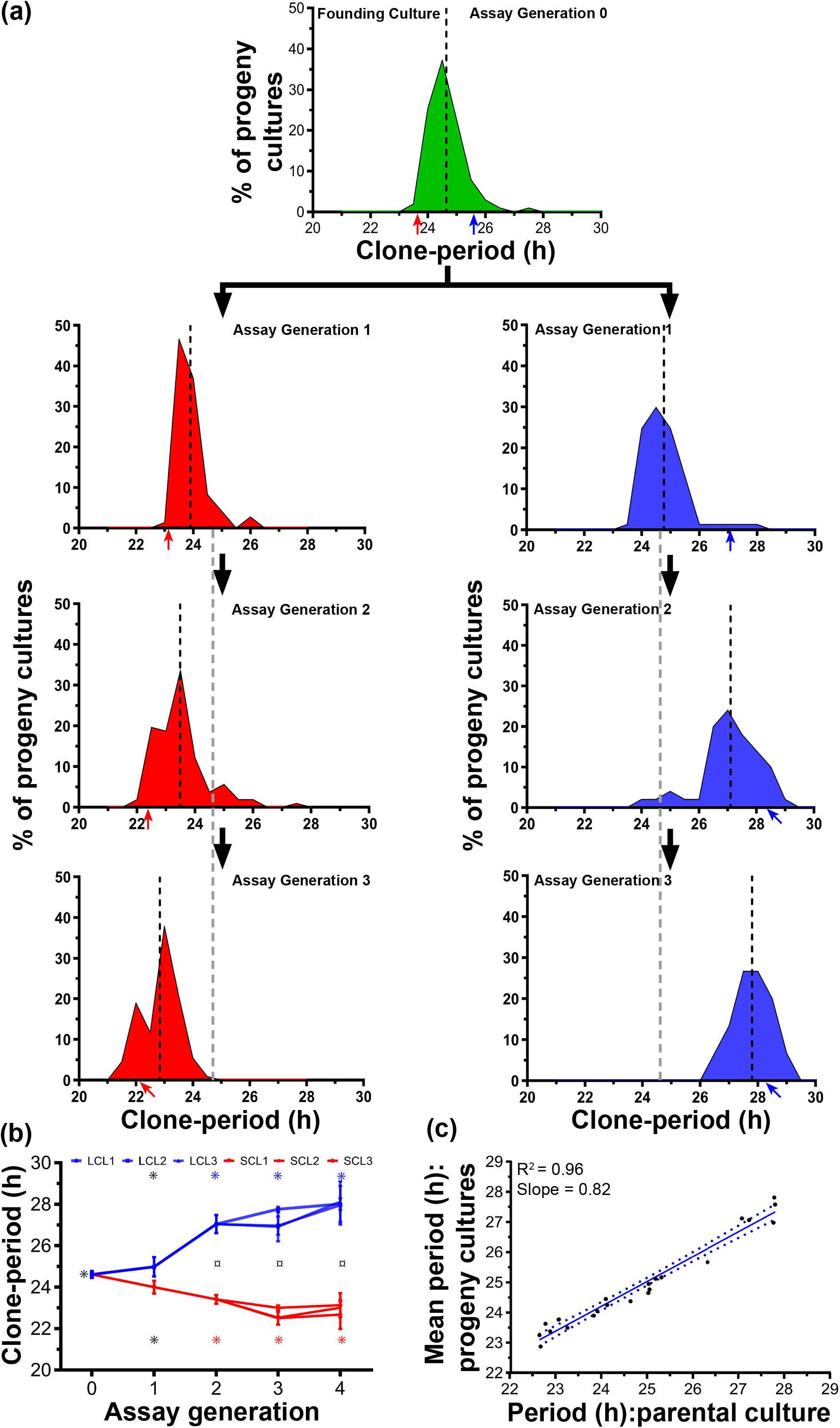
Clonal period-heterogeneity is not stochastically driven but largely inherited. **(a)** Divergence of the period-distributions of short (red) and long (blue) period clones from a common founding culture (green) across multiple assay generations. Dashed black lines depict the mean of respective period distributions. The grey dashed line extended from assay generation-1 depicts mean period of the founding culture (assay generation 0) for visual assessment of the period divergence. Red (short period clone) and blue (long period clone) arrows indicate the means periods of representative clones selected for the successive generation **(b)** Divergence of the mean period among three representative clonal lines each for long-period clonal line (LCL) 1-3 and short-period clonal line (SCL) 1-3. Error bars are SD (n = 3-5 experiments). ¤ indicates that the period of all three SCLs differs significantly from all three LCLs for the given assay generation. Asterisks (*) on top represent LCLs and those at the bottom represent SCLs. Asterisks of different colours indicate that the period of the three clones in that generation is significantly different (*p* < 0.001; n = 3-5) from the periods in other generations, while those with the same colour do not differ significantly. For example, the periods of SCLs 1-3 in assay generation-2 differ significantly from their periods in assay generation-1 and from the founding culture in assay generation-0, but not from assay generations 3 and 4. **(c)** Regression of progeny cultures’ periods on mean periods of their parental cultures’ periods as a proxy-estimate of heritability. Each data point is an average of 3-5 experiments. Blue solid line is the linear regression fit with its 95% CI (blue dotted line).

Interestingly, by repeating this protocol for several assay-generations we observed a directional divergence of the progeny period-distributions on either side of the ‘founding culture’s’ distribution (Figure 1a). The mean circadian periods of progeny cultures in every assay-generation were always very similar to those of their parental cultures (Figure 1a). Over the course of the selection protocol, the periods of short and long period clonal lines (SCL and LCL) significantly diverged from each other and from the ‘founding culture’. The periods of both SCLs and LCLs diverged significantly by assay generation-2 and this divergence reached saturation as the periods did not diverge further (*p* < 0.001; ANOVA followed by Unequal N HSD; Figure 1b). At assay generation 4, the circadian periods of LCLs were ∼3.4 h longer, and those of SCLs were ∼1.7 shorter than the ‘founding culture’s’ period (Figure 1b; Supplementary Figure S2).

As a measure of period-heritability, we regressed mean periods of the progeny cultures on parental cultures and observed that parental period is a very good predictor of the mean progeny period (R^2^ = 0.96; Figure 1c). In addition, even though the divergence of period saturated over the last three assay generations (Figure 1b), we still observe a distribution of periods even after three assay generations. We reason that this distribution might be due to non-heritable stochasticity. Taken together, our results suggest that clonal period-heterogeneity might partly be stochastically driven but has a significant heritable component.

### Inheritance of differential gene expression levels might underlie period heritability

We further aimed to explore the likely basis for heritability of the circadian period underlying heterogeneity. During the course of our experiments, we observed that the short and long period clones consistently exhibited low and high bioluminescence intensities/levels respectively (Supplementary Figure S2a). This encouraged us to test correlation of the period with circadian rhythm parameters such as amplitude, damping rate and bioluminescence intensity.

We observed a positive correlation of bioluminescence intensity (Pearson r = 0.65, *p* < 0.0001) with clone-period; the correlation of relative amplitude with period was negative but not significant (Pearson r = −0.08, *p* = 0.19) and damping rate was not significantly correlated with period either (Spearman r = 0.26, *p* = 0.06; Supplementary Figure S3a). We reasoned that mean bioluminescence intensity can, in-principle serve as a proxy for the average expression level of the underlying gene (*BMAL1* in this case) and hypothesized that clonal inheritance of average gene expression might underlie the observed period heritability. This was further supported by the observation that parental bioluminescence intensity was the best predictor of the respective progeny values (R^2^ = 0.76; Figure 2a), while relative amplitude (R^2^ = 0.04; Supplementary Figure S3b) and damping rate (R^2^ = 0.40; Supplementary Figure S3c) were only poor predictors.

**Figure 2:**
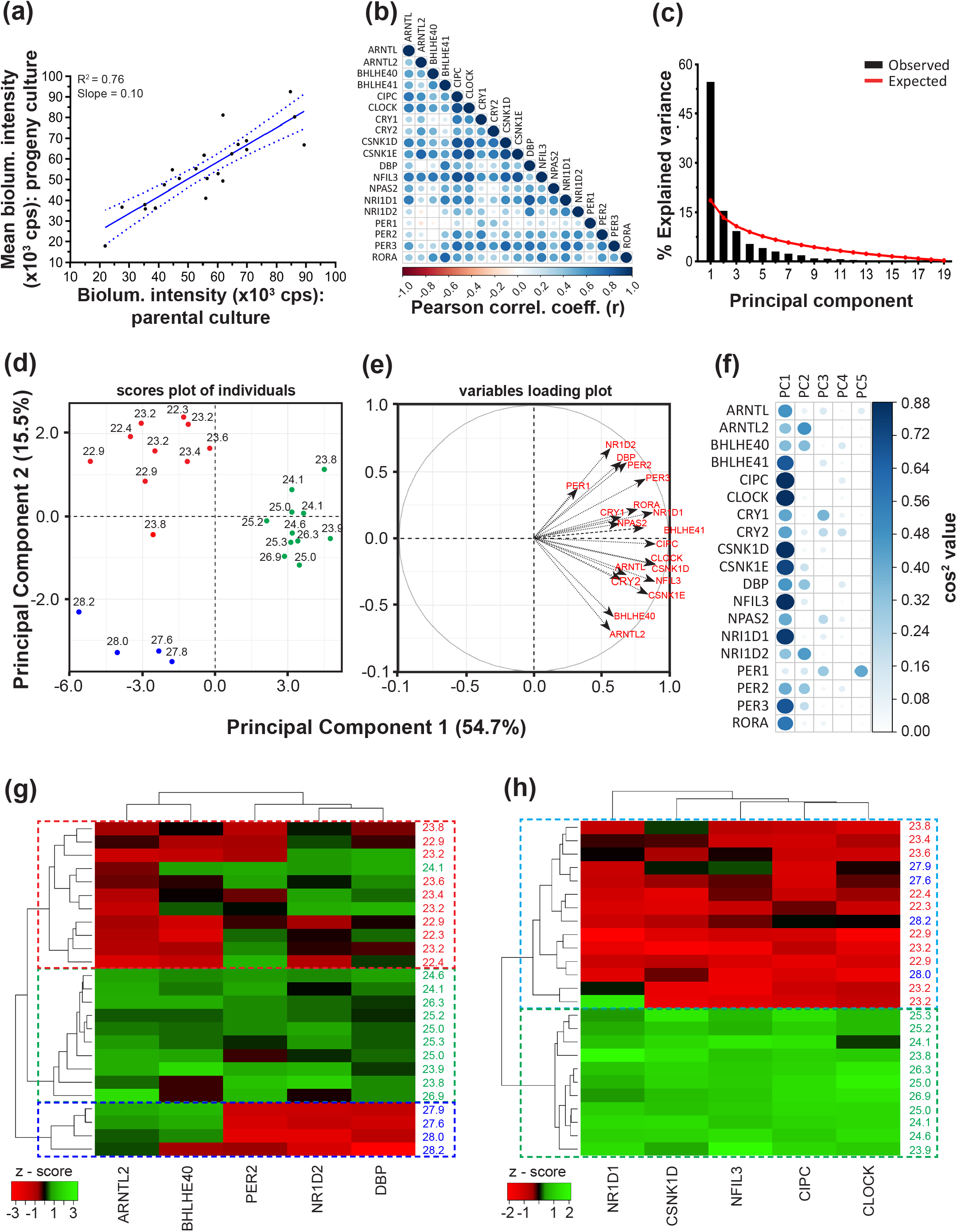
Inheritance of clock-gene expression patterns might govern clonal period-heterogeneity. **(a)** Linear regression of progeny mean bioluminescence intensity on parental values suggests a strong heritability of mean bioluminescence intensity (R^2^ = 0.76). Each data point is an average of 3-5 experiments. Blue solid line is the linear regression fit with its 95% CI (blue dotted line). **(b)** Cross-correlation of average expression values between the 19 analysed genes across all 25 clones indicates a high degree of inter-gene correlation. The colour and size of the circles represent the strength of correlation (Pearson *r*). **(c)** Scree plot depicting the percentage of variance explained by the 19 principal components (black bars) and the expected values based on the Broken-Stick model (red line). **(d)** Factor map of individual clones plotted across the principal components (PCs) 1 and 2 reveals that the first two PCs clusters the clones in three clusters of short (red), intermediate (green) and long (blue) period clones. **(e)** Correlation circle depicting the loading of 19 genes across PCs 1 and 2. **(f)** Cos^2^ values (squared loadings as a measure of the quality of representation of a gene on a PC) of the 19 genes for PCs 1-5. The colour and size of circles represent the magnitude of Cos^2^ value. **(g)** Hierarchical clustering based on the expression of 5 genes selected from PC2. With the exception of one clone, all others clustered into 3 groups of short, intermediate and long clones (red, green and blue dashed rectangles respectively). **(h)** Hierarchical clustering based on the expression of 5 genes from PC1 resulted in 2 clusters – i) intermediate period (green dashed rectangle) and ii) short and long period (blue dashed rectangle). The colour coding of clones in (g) and (h) is the same as in (d).

### Differential expression of E-Box associated factors may govern clonal period-heterogeneity

To test, whether gene expression correlates with clonal period-heterogeneity, we used the NanoString multiplex technology to measure the average expression levels of 20 clock and clock-associated genes (Supplementary Table S1) across our panel of 25 clones. Not surprisingly, we observed a high degree of cross-correlation in expression of the measured genes (Figure 2b) likely due to the high interconnectivity in the circadian clock molecular loop. We subjected the dataset to Principal Component Analysis (PCA) aiming to extract the major features/genes that might underlie (or is a major contributor to) clonal-period heterogeneity. Based on the Broken-Stick model (Jolliffe, 2011), we retained the first two Principal Components (PCs) which collectively explained 70.2% of the circadian period variance (Figure 2c). Interestingly, the first two PCs also clustered the panel of clones into three categories of short (22.3-23h), intermediate (23.8-26.9h) and long (27.6-28.2h) period clones (Figure 2d). PC1 clustered the clones into two groups: i) intermediate periods and ii) the rest including both short and long period clones (non-intermediate). In contrast, PC2 appeared to be important for the three observed clusters (Figure 2d). Based on the cos^2^ values (a measure of the quality of representation of the genes on a PC; Figure 2e-f) and contributions of genes to PC2 (Supplementary Figure S4), we shortlisted the top 25% of the candidate genes (*ARNTL2*, *BHLHE40*, *DBP*, *NR1D2*, *PER2*) that we hypothesized to largely account for the period-variation.

We implemented hierarchical clustering on our dataset based on expression of the five shortlisted candidate genes and observed that clustering of clones was similar (with one exception) to that by the first two PCs (Figure 2g). The amalgamation schedule suggested a possibility of three clusters (red, blue and green dashed-rectangles, Figure 2g) which was also in agreement with the optimal cluster number reported by five different indices (Supplementary Figure S5).

Clustering-based heat map revealed that the expression of *ARNTL2* and *BHLHE40* correlated positively with the circadian period, *DBP* and *NR1D2* correlated negatively, while *PER2* exhibited a clear trend (Figure 2g, Supplementary Figure S6a). As a control measure, we also similarly shortlisted top 25% genes from PC1 (*NR1D1*, *CLOCK*, *CSNK1D*, *CIPC* and *NFIL3*) and, as expected, we observed that these genes were not sufficient to discriminate the short and long periods thereby resulting in only two clusters – intermediate and non-intermediate (Figure 2h). Interestingly, all five genes from PC1 have higher expression in ‘intermediate’ period clones and their expression reduces as the period deviates from ‘intermediate’ (Figure 2h, Supplementary Figure S6b). Thus, we reasoned that changes in expression of PC2 genes are likely to drive period heterogeneity while those from PC1 are likely to be a consequence of period heterogeneity.

We hypothesized that if differences in expression of the shortlisted PC2 genes governs period heterogeneity, then depletion of these genes should result in large period change while depletion of those from PC1 should not have a significant effect on period. Specifically, based on their expression patterns (Figure 2g) knockdown of *ARNTL2* and *BHLHE40* should shorten the circadian period while *DBP* and *NR1D2* knockdown should result in period lengthening. To test this, we used RNAi mediated silencing to individually knockdown the shortlisted genes in 3-short, 2-intermediate and 3-long period clones (based on clustering in Figure 2g) and studied the effect on circadian period. Indeed, we observed that knockdown of *NR1D2* resulted in significant period lengthening across all clones while *BHLHE40* and *ARNTL2* knockdown resulted in significant period shortening (Mixed model ANOVA followed by Tukey’s HSD; *p* < 0.00001; Figure 3a-b). *NR1D2* knockdown had the largest effect on period, significantly higher compared to all other genes across both the PCs; followed by *BHLHE40* that was similar to *ARNTL2* and had a significantly higher effect on period compared to all other genes. Knockdown of none of the other genes across both PCs resulted in a period change significantly differing either from zero (one sample t test, *p* > 0.05) or from each other (Mixed model ANOVA followed by Tukey’s HSD; *p* > 0.05; Figure 3a-b). Accordingly, we observed that the average absolute period change upon knockdown of PC2 genes was significantly higher than that by PC1 genes (Figure 3c).

**Figure 3:**
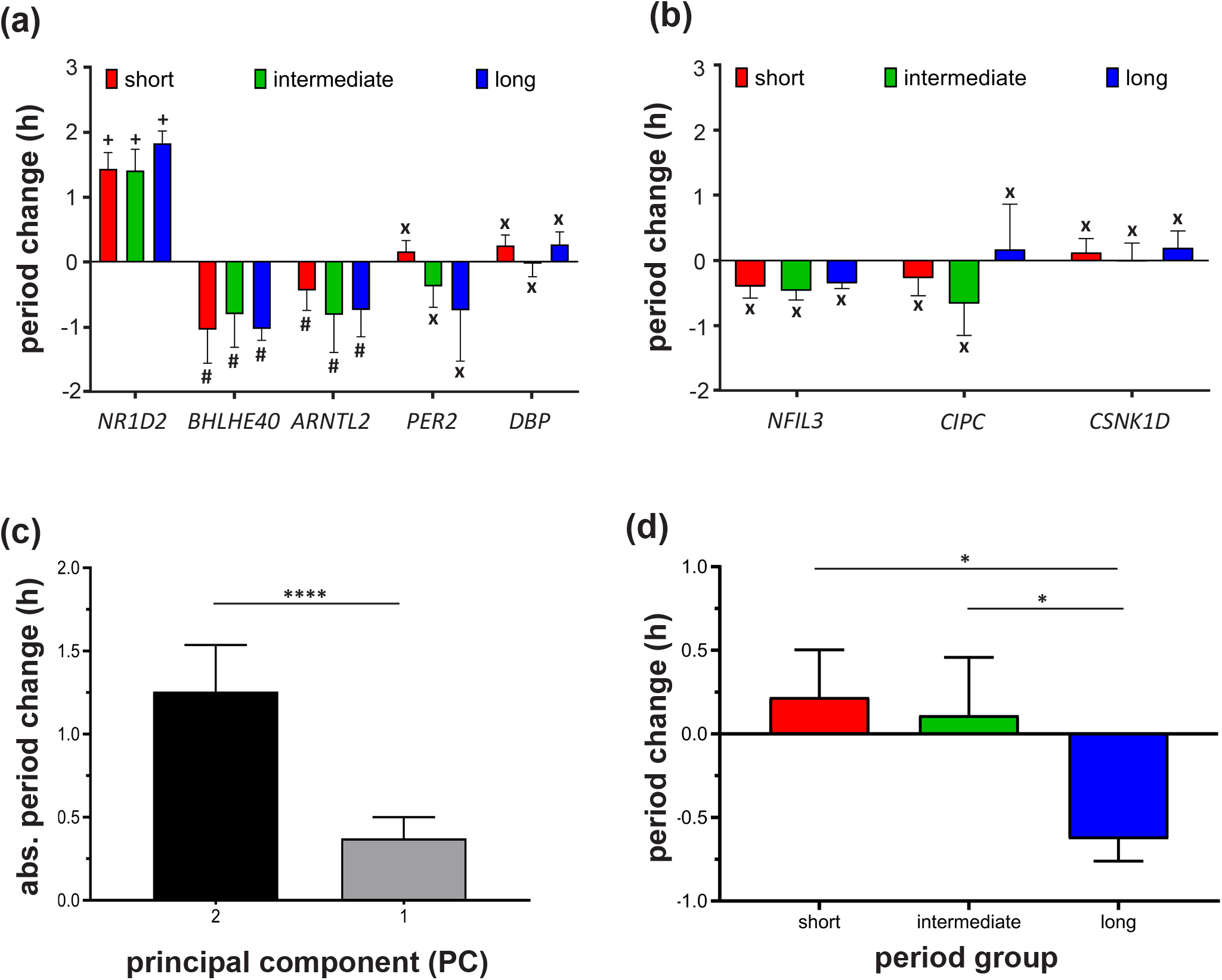
Epigenetically regulated expression of E-Box associated factors may govern clonal period-heterogeneity. Period change (compared to non-silenced control) upon knockdown of the **(a)** five PC2 genes and **(b)** three PC1 genes for the short (red), intermediate (green) and long (blue) period clones. Bars with different symbols indicate significant differences (*p* < 0.05) while bars with same symbols are not significantly different from each other (Mixed model ANOVA followed by Tukey’s HSD). **(c)** Averaged absolute period change across all clones upon knockdown of genes from PC2 (black) and PC1 (grey). **(d)** Period change (compared to vehicle control) upon treatment of short (red), intermediate (green) and long (blue) period clones with HDAC (Histone Deacetylase) inhibitor SAHA (1.6 µM). For all panels in this figure, n = 3-4 experiments and error bars are SD. *: *p* < 0.05; ****: *p* < 0.0001.

Taken together, these results suggest that differential expression of *NR1D2*, *BHLHE40* and *ARNTL2* likely underlies heterogeneity in clonal circadian period.

### Epigenetic regulation might underlie altered gene expression patterns associated with clonal period-heterogeneity

Having observed that clonal period-heterogeneity is associated with altered gene expression patterns, we next asked ‘what causes such altered expression among clonal cells?’ We ruled out the possibility of random mutation accumulation (see Discussion) and hypothesized that epigenetic variation might account for the observed differences in gene expression patterns among clonal lines. As a preliminary test, we treated all 25 clonal cell populations in our panel with the commonly used epigenetic modifier Suberoylanilide Hydroxamic Acid (SAHA) and studied the effect of the treatment on clone period. SAHA is a Class I and Class II Histone Deacetylase Inhibitor which upregulates gene expression by multiple mechanisms (Marks *et al*., 2008). We reasoned that if reduction in expression of the identified subset of genes across our clonal panel is due to epigenetic suppression (in this case, acetylation status), treatment with SAHA should upregulate the expression of these genes thereby lengthening and shortening the circadian period in short and/or long period clones respectively.

Interestingly, we observed that treatment with SAHA differentially influenced the short, intermediate and long period clones. SAHA treatment resulted in a significant period shortening in the long period clones (ANOVA followed by Tukey’s HSD, *p* < 0.05; Figure 3d) whereas, the magnitude of period change in short and intermediate period clones did not differ from each other (ANOVA followed by Tukey’s HSD, *p* = 0.85) or from zero (one sample t test, *p* > 0.05; Figure 3d).

Although the possible reasons for the differential effects of SAHA treatment on short and long period clones will be discussed later, this provides preliminary evidence suggesting that epigenetically regulated gene expression differences might underlie clonal period-heterogeneity.

## DISCUSSION

We used human U-2 OS cells to investigate whether period-heterogeneity in circadian clock network stems from intrinsic/extrinsic non-heritable stochasticity or whether it has a heritable component. We employed a laboratory selection protocol to select for clonal cell populations exhibiting short and long circadian periods through which we derived a panel of 25 clonal cell populations exhibiting circadian periods between 22 h to 28 h.

We observed that parental clones always produced progeny with mean periods closely resembling the former thus resulting in a directional response (divergence of short and long period clones from the founding culture) to our selection protocol (Figure 1a). Consistently, the period of parental culture was a very good predictor (R^2^ = 0.96) of the progeny’s mean period (Figure 1c). Taken together, these results suggest that clonal period-heterogeneity is unlikely to be stochastically driven and has a strong heritable component. This raises an interesting question: why would natural selection favour the evolution of heritable mechanisms to drive period-heterogeneity over entirely stochastically driven heterogeneity? We hypothesize that, although period heterogeneity can be functionally beneficial (Jagota *et al*., 2000; Schaap *et al*., 2003; Gonze *et al*., 2005; Bernard *et al*., 2007; Inagaki *et al*., 2007; VanderLeest *et al*., 2007; Gu *et al*., 2016, 2019), very large heterogeneity can negatively influence clock functionality as well (Gonze *et al*., 2005; Bernard *et al*., 2007; Gu *et al*., 2016). Stochastic mechanisms can potentially lead to very large variation in inter-cellular/oscillator period which would be detrimental, whereas heritable mechanisms may impose phenotypic constraints (Wagner, 2011) within which period-heterogeneity can be maintained and thus being favoured by natural selection.

Over the course of our experiments, we observed that long-period clones often exhibited higher bioluminescence intensity compared to the short-period clones (Supplementary Figure S2a) and further analysis revealed that parent bioluminescence intensity was a good predictor of progeny bioluminescence intensity but this was not the case for either relative amplitude or the damping rate (Figure 2a, Supplementary Figure S3). We reasoned that bioluminescence intensity could serve as a proxy measure for *BMAL1* expression and thus hypothesised that period heritability is likely to be due to the inheritance of gene expression levels from parental clones. To further explore this, we measured average expression of 20 circadian clock and clock-associated genes (Supplementary Table S1) across all 25 clones in our panel. By employing principal component analysis, we identified five candidate genes (*ARNTL2*, *BHLHE40*, *DBP*, *NR1D2* and *PER2*) that grouped the clones into three distinct clusters – short, intermediate and long periods (Figure 2c-g, Supplementary Figures S4-5). Furthermore, we observed that knockdown of three of the shortlisted candidates - *NR1D2*, *BHLHE40* and *ARNTL2* had the largest influence on period across while other genes including those from PC1 had little or no effect on period change (Figure 3a-c). It is noticeable that individual knockdown of the genes resulted in small magnitude period changes that cannot entirely account or period differences between the short and long period clones (Figure 3a). These results suggest that that clonal period-heritability is a multi-gene trait involving a consortium of multiple medium-effect genes. Notably, all three above-mentioned genes are transcription factors that are either regulated by and/or act on E-boxes and are part of both the core and auxiliary molecular clock loops (Ikeda *et al*., 2000; Okamura *et al*., 2002; Kawamoto *et al*., 2004; Guillaumond *et al*., 2005; Nakashima *et al*., 2008; Sasaki *et al*., 2009; Takahashi, 2017). This reinforces the idea that while persistence of circadian oscillation requires a functional core clock loop involving negative feedback by the PER-CRY family, modulation of clock period might be governed by interaction between multiple loops coupled by E-box associated transcription factors (Zhang and Kay, 2010; Relógio *et al*., 2011). Another notable gene that our analysis revealed happens to be one of the relatively less studied circadian clock genes *ARNTL2* (*BMAL2*). While *ARNTL2* is a functional paralog of the core clock gene *ARNTL1* (*BMAL1*), its precise role in the clock loop remains largely elusive (Ikeda *et al*., 2000; Sasaki *et al*., 2009; Shi *et al*., 2010) thus highlighting a potential role of *ARNTL2* in circadian period-modulation, which awaits further exploration.

Intriguingly, in contrast to the above-discussed genes, we find another category among the assayed genes that exhibit an inverted-U shaped relationship with period. The expression of these genes (*NR1D1*, *CSNK1D*, *NFIL3*, *CLOCK*, *CIPC*) is high in clones with intermediate periods (23.8-26.9h) and is drastically reduced in clones with periods deviating from the intermediate range (Figure 3g). Furthermore, our knockdown studies also confirm that expression patterns of these genes are not causal but likely to be a response/consequence to period variation (Figure 3b). Such inverted-U shaped responses (Hormesis) is observed in various biological systems and is regarded as a regulatory/homeostatic mechanism to prevent very large deviations of cellular/organismal phenotypes from their optimal range (Calabrese and Baldwin, 2001; Baldi and Bucherelli, 2005; Zhang *et al*., 2008). As discussed earlier, since a higher degree of period-heterogeneity can be detrimental to the circadian clock network, we hypothesize that while there are mechanisms within the clock circuitry that promote period-heterogeneity, the network might also harbour hormesis-based mechanisms which impose constraints on the range of period that the circadian clock can exhibit (Baldi and Bucherelli, 2005; Zhang *et al*., 2008). Such mechanisms may also explain why we observe a saturation of period divergence after assay generation 2 (Figure 1b).

While evidence thus far strongly suggests that clonal period-heterogeneity is driven by differences in clock gene expressions, we then asked ‘what is the source of these expression differences?’ One possibility is that the short and long period clones might have accumulated random mutations resulting in period change and subsequently selected by us. However, we reason that this is highly unlikely because – a) With a mutation rate of ∼2.5 x 10^-8^/nucleotide in human cells (Nachman and Crowell, 2000), the probability of occurrence of at least two kinds of mutations within a small fraction of the genome (comprising clock genes) driving short and long periods is extremely low. b) We see significant trends in expression of the same subset of genes across both short and long period clones (Figure 2g; Supplementary Figure S6). This presupposes that mutations driving short and long periods have occurred within the same genes, which further drastically reduces the probability that the observed period differences stem from random mutations. c) Even if the mutation rate is higher than we estimate, the saturation of divergence in period over the last three assay generations (Figure 1b) cannot be entirely accounted by mutations since the periods could continue to diverge due to further accumulation of mutations. Therefore, we argue that the observed period-heterogeneity is unlikely to be due to random mutations, which leaves us with another alternative – epimutations. Epimutations are heritable changes in expression of genes and are not associated with DNA mutations. Epimutations are often associated with changes in methylation states of genes or other heritable chromatin modifications (Holliday, 2006). The rates of epimutations are observed to be order of magnitude higher than DNA mutation rates (Van Der Graaf *et al*., 2015) and successfully explains phenotypic heterogeneity in many life forms including clonal populations (Kaufmann *et al*., 2007; Stockholm *et al*., 2007; Neildez-Nguyen *et al*., 2008; Taudt *et al*., 2016; Springer and Schmitz, 2017). Therefore, we hypothesized that epimutations-driven gene expression differences may underlie clonal heterogeneity in circadian period. As a preliminary test of this hypothesis, we studied the effect of a Histone Deacetylase Inhibitor Suberoylanilide Hydroxamic Acid (SAHA) treatment on the circadian period across our clones. Interestingly, we find that treatment with SAHA significantly shortens (albeit by a small magnitude) the period in long-period clones with little or no effect on the short and intermediate clones (Figure 3d). The small magnitude effect of SAHA treatment might be due to one or all of the following reasons. SAHA is broad spectrum Histone Deacetylase (HDAC) inhibitor and promotes upregulation of genes by acetylation, whereas other epigenetic mechanisms that might contribute to the gene expression in our clones are not targeted by this treatment. Alternatively, off-target effects of SAHA might also upregulate other genes that in turn negatively influence the change in period. In addition, as discussed previously, if period heterogeneity is indeed a multi-gene trait relying on combined upregulation and downregulation of two or more genes, mere treatment with epigenetic modifiers that leads to genome-wide changes in gene expression may not be a good strategy. Nevertheless, the differential effects of SAHA on short and long period clones is promising and provides preliminary support to the idea that epigenetic modulation of gene expression might underlie clonal period-heterogeneity. Future targeted studies along these lines may shed more light on this aspect.

In conclusion, our study reports that the heterogeneity in periods observed within circadian clock networks in mammals is not stochastically driven but has a heritable basis and that this is likely to be a multi-gene trait. We identified that differential regulation of three E-box associated transcription factors might govern period-heterogeneity in circadian clock networks and provide preliminary evidence that epigenetically regulated gene expression differences may underlie clonal period-heterogeneity. In addition, we also observed a subset of genes that exhibit which we hypothesize are part of homeostatic mechanisms that may constrain circadian clocks from deviating largely from their optimal period range. Future studies will help further explore the phenomenon of period-heterogeneity and its regulation.

## MATERIALS AND METHODS

### Clone selection protocol

All clones used in this study were U-2 OS cells (human, ATCC # HTB-96) stably expressing firefly luciferase from a 0.9-kb *BMAL1* promoter (Maier *et al*., 2009), cultured and maintained in DMEM containing 10% fetal bovine serum, antibiotics (100 U/ml penicillin and 100 μg/ml streptomycin). See Supplementary Figure S1 for a pictorial description of the selection protocol. Briefly, cells from ‘founding culture’ expressing a circadian period of 24.6 ± 0.16 h (mean ± SD) were plated as single-cell clones in 96-well ‘parent plates’ and grown to confluency. Upon reaching confluency, an ‘assay plate’ was established for every ‘parent plate’ by splitting cells. The period of bioluminescence rhythms from cells in ‘assay plates’ was recorded (see below for recording protocol) and clones exhibiting short or long periods (tails of the period-distribution) were selected. Bioluminescence rhythms of every clone was recorded 2-3 times and only clones that consistently exhibited shot/long periods were selected. Following the selection of clones, corresponding clones from the ‘parent plate’ were single-cell cloned in 96-well plates, and the procedure was repeated for four assay generations by selecting short and long period clones every generation.

### Bioluminescence recording

Cells were plated in white 96-well plate at a density of 20×10^3^ cells/well and after 72 hours, cells were synchronized with dexamethasone (1 μM) for 30 minutes, washed twice with PBS and cultured in Phenol-Red-free DMEM containing 10% fetal bovine serum, antibiotics (100 U/ml penicillin and 100 μg/ml streptomycin) and 250 μM D-luciferin (Biothema, Darmstadt, Germany). Bioluminescence was recorded at 37°C in a 96-well plate luminescence counter (TopCount, PerkinElmer, Rodgau, Germany) for up to 7-days. ChronoStar software (Maier *et al*., *in press*) was used for data analysis and estimation of rhythms parameters including period, decay constant (damping), relative amplitude and average bioluminescence (MESOR) of the oscillation as described previously (Abraham *et al*., 2010).

### RNA preparation and NanoString based gene expression analysis

Five days before the RNA extraction, cells were plated at a density of ∼20×10^3^ cells/well in 24-well plate with DMEM containing 10% fetal bovine serum, antibiotics (100 U/ml penicillin and 100 μg/ml streptomycin). Since we intended to measure average gene expression levels, the culture medium was not replaced for five days to prevent accidental synchronization of cells. On day-5 the medium was removed, 100 μl/well iScript™ RT-qPCR Sample Preparation Reagent (Biorad) was added on top of the cell-layer and incubated at 37 ^0^C for 5 min. 3μl of the sample was withdrawn without disturbing the cell-layer and used for further downstream analysis as per manufacturer’s instructions.

A previous study of ours combined whole-genome transcriptomics with machine learning and identified genes that could serve as reliable circadian time-telling markers (Wittenbrink *et al*., 2018). Based on this, we designed a 24-plex NanoString probe panel comprising 20 circadian clock and clock associated genes and 4 housekeeping genes (Supplementary Table S1). The custom-designed probes included a 3′-end biotinylated capture probe and a 5′-fluorescence-barcoded reporter probe for each gene target. Hybridization of probes and gene expression-count reading was according to the manufacturer’s instructions. Raw expression data was acquired by a NanoString nCounter Digital Analyzer (NanoString Technologies), QC processed and analysed by nSolver^TM^. QC analysis flagged reads from one (*CIART*) of the 24 genes in the panel as unsuitable for analysis and was not considered. Data normalization involved three steps: (a) normalization by the arithmetic mean of the positive spike-in controls, (b) subtraction of the mean of the negative controls, and (c) normalization by the geometric mean of the four housekeeping genes.

### Principal Component Analysis and Clustering

Log2-transformed gene expression data were first subjected to Bartlett’s Test of Sphericity to validate its adequacy for Principal Component Analysis (PCA) following which correlation-based PCA was implemented in R (R Core Development Team, 2013) using factoextra and FactoMineR packages (Kassambara, 2016). Broken-Stick model (Jolliffe, 2011) was used to determine the number of retainable Principal Components (PCs). Determining the optimal cluster-number is often a complication in unsupervised exploratory data analysis. Unlike many studies in biology that employ PCA to identify genes based on expression differences between known cell-types (which can be used to estimate the optimal number of clusters), our study employs a panel of clones with a continuous distribution of phenotypes (period) and thus cannot be categorized trivially. Hence, we adopted two schemes for optimal cluster-number determination. (a) For agglomerative hierarchical clustering, we assessed the agglomeration schedule to identify the possible number of clusters (Yim and Ramdeen, 2015). (b) In addition, we also performed k-means clustering for different values of cluster (k = 1-10) and used 5 different indexes – ‘silhouette method’ (Rousseeuw, 1987), ‘elbow method’ (Thorndike, 1953), ‘gap-statistic’ (Tibshirani *et al*., 2001), ‘Calinski-Harabasz criterion value (variance-ratio method)’ (Caliñski and Harabasz, 1974) and Bayesian Information Criterion (BIC; Fraley and Raftery, 2002) to assess the optimal cluster-number. We selected the optimal cluster number based on agreement between (a) and (b). Heatmapper (Babicki *et al*., 2016) and ‘dendextend’ (Galili, 2015) were used for hierarchical clustering analysis based on ‘euclidean-distance’ and ‘complete-linkage’ measures (D, 2005). ‘Nbclust’ (Charrad *et al*., 2014) and ‘mclust’ (Scrucca *et al*., 2016) were used for k-means based clustering analysis while for all other statistical analysis and graphing was performed using R and Prism version 8.00 for Windows (GraphPad Software, La Jolla California USA, www.graphpad.com).

### RNAi mediated gene knockdown

The GIPZ microRNA-adapted shRNA constructs used for the study were purchased from Open Biosystems and packaged into lentiviral vectors in HEK293T cells in a 96-well plate format (Maier *et al*., 2009). Virus-containing supernatants were then filtered and reporter cells (clonal cell populations used in the study) were transduced with 150 μL of the filtrate containing 8 ng/μL protamine sulfate. After at least 24h, the filtrate was replaced with fresh medium containing puromycin (10 μg/mL). After 3 days, the transduced reporter cells were synchronized and bioluminescence rhythms were recorded as described above.

### SAHA treatment and dose response analysis

10^3^cells were plated in a 96-well plate on day-0. After 24h, the culture media was replaced with media containing 1.6 µM Suberanilo Hydroxamic Acid (SAHA) or DMSO vehicle control. The drug was replaced every day for three consecutive days. On day-4, the cells were rinsed thrice with PBS and fresh (no drug) culture media was added. The cells were untreated for next 48h and bioluminescence rhythms were recorded from day-6.

The above-described protocol was followed for estimating the IC50 value as well. Cells were treated with varying concentrations (0-100 µM) of SAHA from day-1, and cell proliferation was assayed on day-6 using the Vybrant® MTT Cell Proliferation Assay Kit (Thermo Fischer Scientific, catalog #V13154) as per manufacturer’s protocol. IC50 was calculated from the resulting dose response curve using Prism version 8.00 for Windows (GraphPad Software, La Jolla California USA, www.graphpad.com; Supplementary Figure S7).

## ACKNOWLEDGEMENTS

The authors thank Sebastian Jäschke and Katja Schellenberg, Astrid Grudziecki and Almut Eisele for their help with single-cell cloning; Hedwig Lammert, Michael Hummel and Barbara Koller for help with NanoString experiments and Lucas Hille for his help with gene-knockdown and epigenetics related experiments. Work in AK’s lab is supported by grants from the Deutsche Forschungsgemeinschaft to SK (SFB740/D2) and AK (TRR186/P17). Alexander von Humboldt-Stiftung supported NKL’s research.

## AUTHOR CONTRIBUTIONS

Conceptualization: NKL, SK and AK; Experiments and data acquisition: NKL and SK. Data curation and formal analysis: NKL and SK; Manuscript preparation: NKL and AK.

