## Supplemental Information for "Heritable gene expression variability governs clonal heterogeneity in circadian period"

#### SUPPLEMENTARY FIGURE LEGENDS

**Supplementary Table S1**

List of clock genes, clock associated genes and 4 housekeeping genes (GAPDH, HPRT, PPIA and PSMB2) used for the study.

**Supplementary Figure S1**

Graphical depiction of the selection protocol adopted for deriving the panel of short and long period clones used in this study. TopCount image courtesy: https://www.perkinelmer.co.uk.

**Supplementary Figure S2**

**(a)** Raw and **(b)** de-trended bioluminescence traces of representative clones from founding culture (green), short period (red) and long period (blue) clonal lines. Time series in (b) is shorter because de-trending results in truncation of data by 24 hours.

**Supplementary Figure S3**

(**a**) Correlations between clone-period and rhythms parameters – relative amplitude (Pearson r = -0.08, $p = 0.19$), bioluminescence intensity (Pearson r = 0.65) and damping rate (Spearman r = 0.26, $p = 0.06$). Bioluminescence intensity and relative amplitude values were normally distributed (KS distance = 0.08, $p > 0.10$ and KS distance= 0.07, $p > 0.10$ respectively) whereas damping rate was not (KS distance = 0.21, $p < 0.0001$). Hence non-parametric Spearman correlation was used for damping rate-clone period correlation. Linear regression of mean progeny values on parental values for **(b)** relative amplitude ($R^2 = 0.04$) and **(c)** damping rate ($R^2$

= 0.40). Each data point is an average of 3-5 experiments. Blue solid line is the linear regression fit with its 95% CI (blue dotted line). ****: $p < 0.0001$

**Supplementary Figure S4**

Contributions of the 19 analysed genes to **(a)** principal component-2 and **(b)** principal component-1. Note that a few genes contribute largely to PC2 whereas PC1 has lower and almost equal contributions from a large number of genes compared to PC2.

**Supplementary Figure S5**

To estimate the optimal number of clusters in our dataset, we measured five different k-mean clustering indexes (average silhouette width, within sum of squares, gap statistic, Calinski-Harabasz value and Bayesian Information Criterion) for k = 1-10 clusters. **(a)**-**(e)** values of the above mentioned indexes across 10 clusters generated by the 5 selected genes selected from principal component-2 and **(f)**-**(j)** represent the same for clusters generated by the 5 selected genes selected from principal component-1. The red dots indicate the optimal cluster number chosen based on the respective index. Details of the indexes used and their interpretation can be found in the respective references (see Materials and Methods). In brief, for all indices except Within-Sum of Squares (WSS), the cluster number resulting in the highest value of the index was considered likely to be the optimal cluster number. For WSS, the cluster number at which the WSS plot forms an elbow-joint (the magnitude of drop in WSS values reduces thereafter) was considered as the likely optimal cluster number. When more than one optimal cluster number is observed, as in (c), (d) and (h), the decision was based on guidelines suggested by the original authors (see Materials and Methods).

**Supplementary Figure S6**

Trends of gene expression across clones exhibiting different circadian periods for the five selected genes from **(a)** Principal Component (PC)-2 and **(b)** Principal Component (PC)-1. r = Pearson correlation coefficient and $R^2$ = goodness of linear regression fit (blue solid lines) to estimate the proportion of variance in clone-period explained by variance in gene expression. *: $p < 0.05$; **: $p < 0.001$; ***: $p < 0.0001$; ****: $p < 0.00001$

**Supplementary Figure S7**

To estimate $IC_{50}$ value for SAHA, cells were treated with varying concentrations of the drug (0 µM - 104 µM) for three days (see Materials and Methods) after which cell proliferation was measured as absorbance at 570 nM using Vybrant® MTT Cell Proliferation Assay Kit (Thermo Fischer Scientific, catalog #V13154). From the resulting dose response curve, $IC_{50}$ was calculated using Prism version 8.00 for Windows (GraphPad Software, La Jolla California USA, www.graphpad.com ; Supplementary Figure S7). Error bars on data points represent SD (n = 3). The solid dashed line is the non-linear regression fit with its 95% CI (dotted line).

### Supplementary Table S1

| Gene | Gene Alias | Accession Number |
|---|---|---|
| ARNTL | BMAL1 | NM_001030272.1 |
| ARNTL2 | BMAL2 | NM_020183.3 |
| BHLHE40 | DEC1 | NM_003670.2 |
| BHLHE41 | DEC2 | NM_030762.2 |
| CIART | CHRONO | NM_144697.2 |
| CIPC | - | NM_033426.2 |
| CLOCK | - | NM_004898.2 |
| CRY1 | - | NM_004075.3 |
| CRY2 | - | NM_001127457.1 |
| CSNK1D | - | NM_001893.3 |
| CSNK1E | - | NM_152221.2 |
| DBP | - | NM_001352.3 |
| NFIL3 | E4BP4 | NM_005384.2 |
| NPAS2 | - | NM_002518.3 |
| NR1D1 | REVERB-α | NM_021724.3 |
| NR1D2 | REVERB-β | NM_001145425.1 |
| PER1 | - | NM_002616.2 |
| PER2 | - | NM_022817.2 |
| PER3 | - | NM_016831.1 |
| RORA | - | NM_134261.2 |
| GAPDH | - | NM_001256799.1 |
| HPRT1 | - | NM_000194.1 |
| PPIA | - | NM_021130.3 |
| PSMB2 | - | NM_002794.3 |

#### Supplementary Figure S1

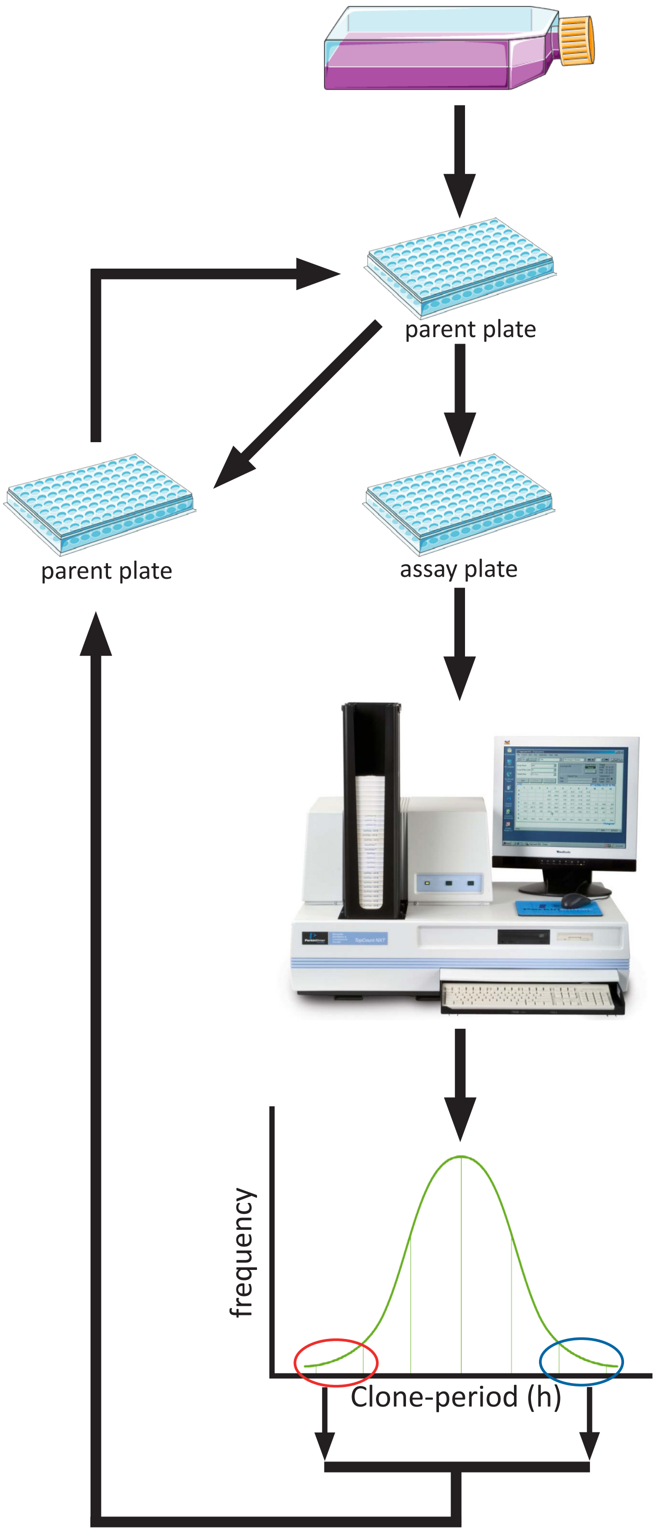


Founding culture

Cells from founding culture were single-cell cloned in a 96-well ‘parent plate’ and cultured till confluency

Upon reaching confluency, an ‘assay plate’ was established by splitting cells from the ‘parent plate’

Cells in ‘assay plate’ were synchronized and their bioluminiscence rhythms were recorded for 5-7 days in TopCount® luminometer

The resulting distribution of periods from all clones were analysed. Multiple clones exhibiting short or long periods were selected for the subsequent generation. The procedure was repeated 2-3 times for every ‘parent plate’ and clones which consistently exhibted short or long period were selected.

The selected clones were then single-cell cloned from the ‘parent plate’, and the protocol was repeated for several generations.

### Supplementary Figure S2

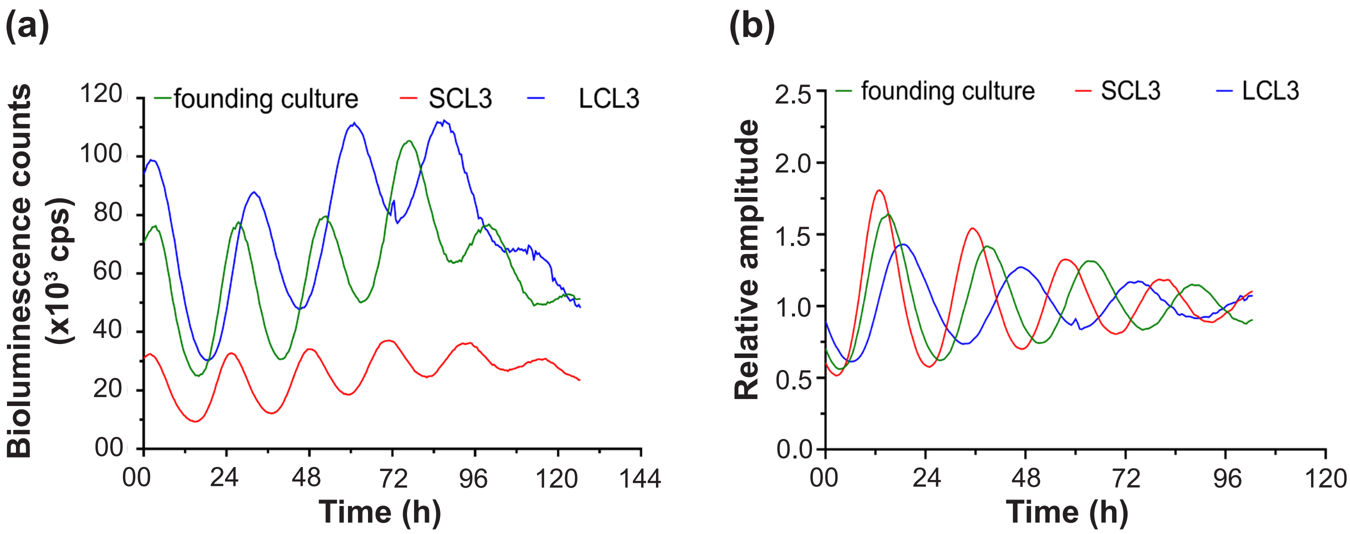

### Supplementary Figure S3

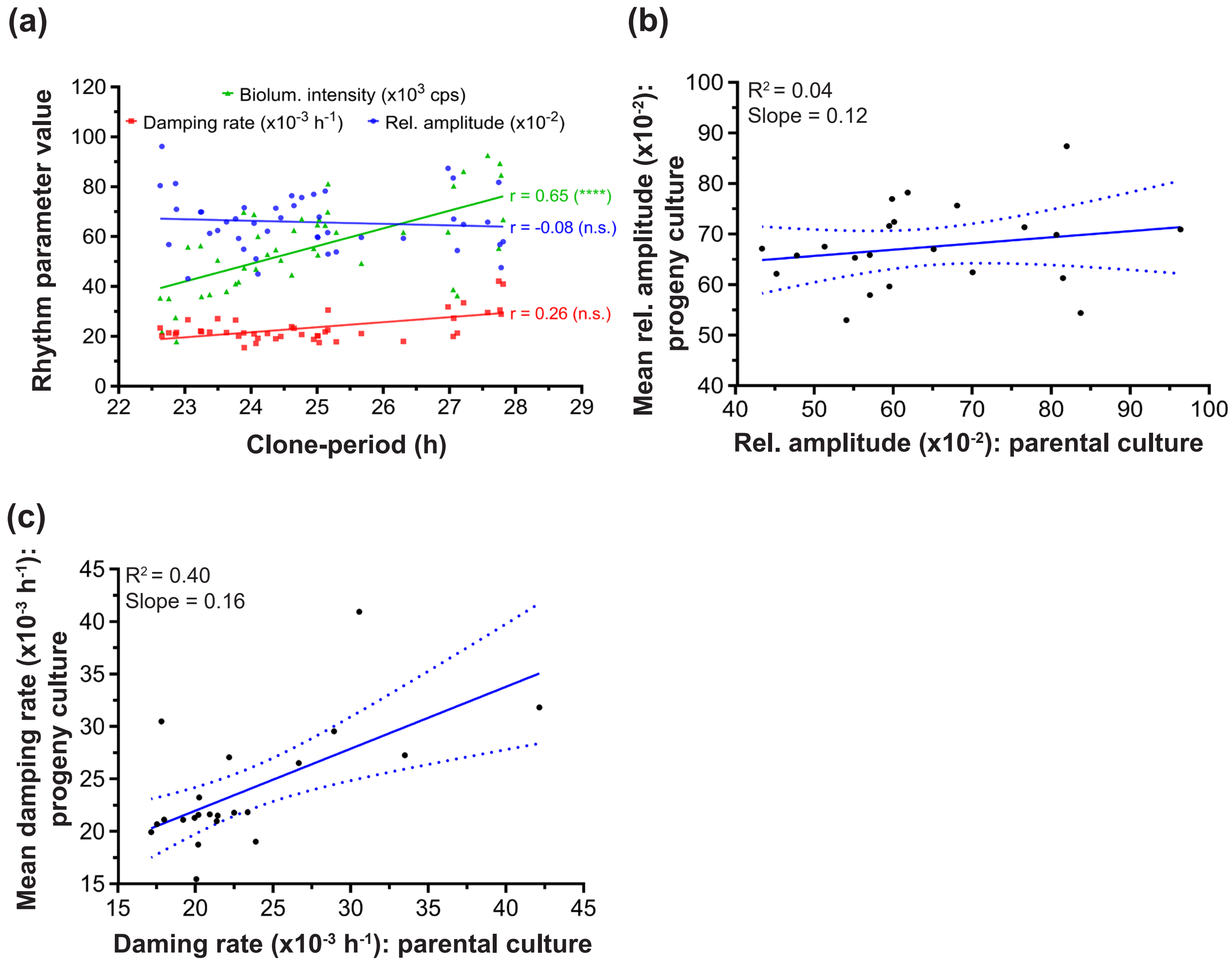

### Supplementary Figure S4

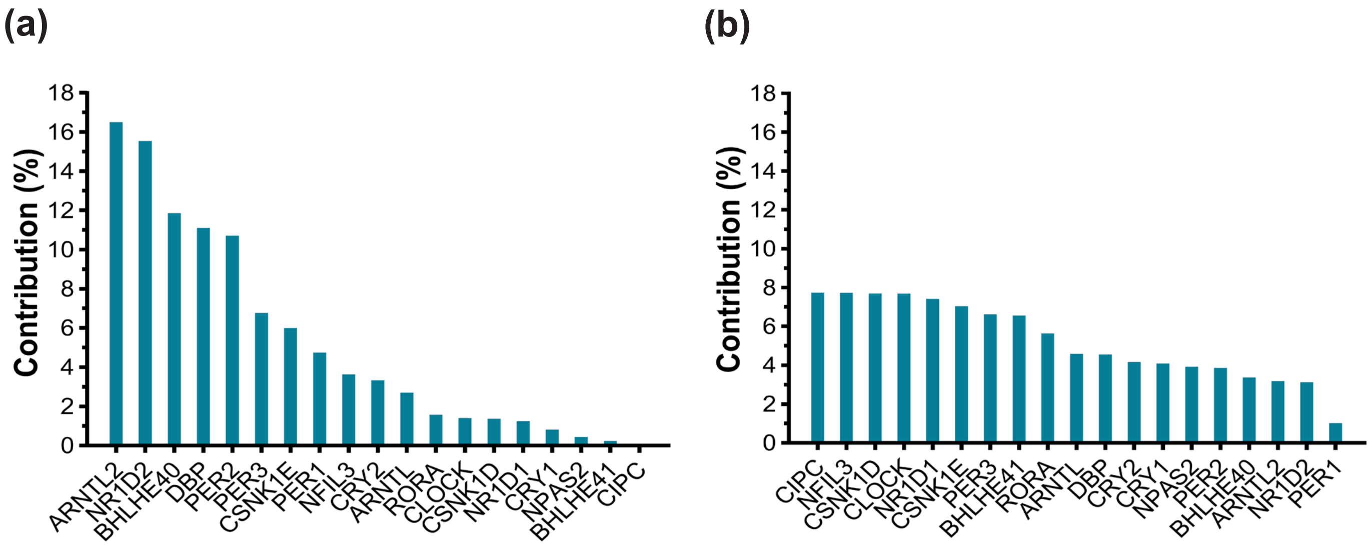

### Supplementary Figure S5

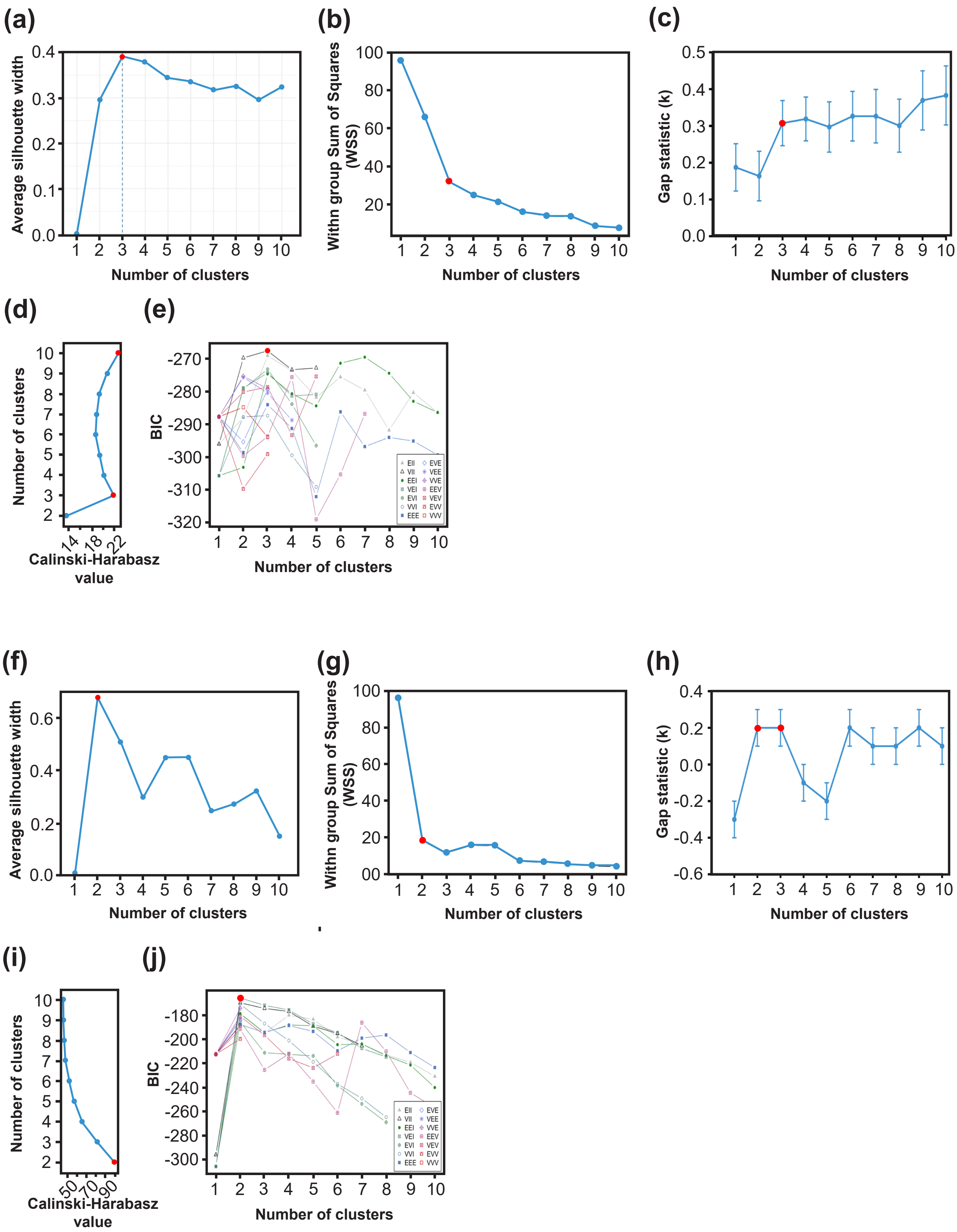

### Supplementary Figure S6

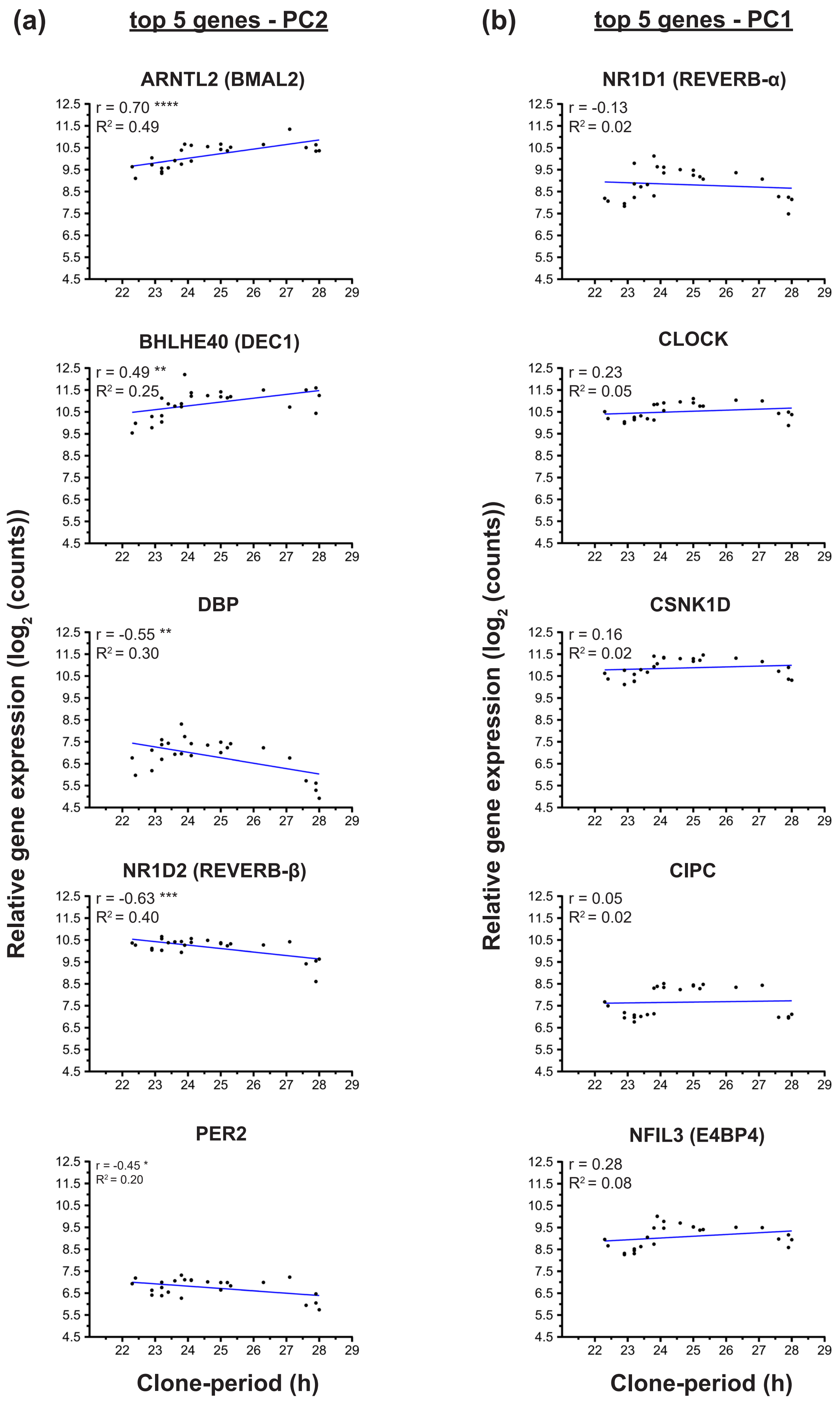

#### Supplementary Figure S7

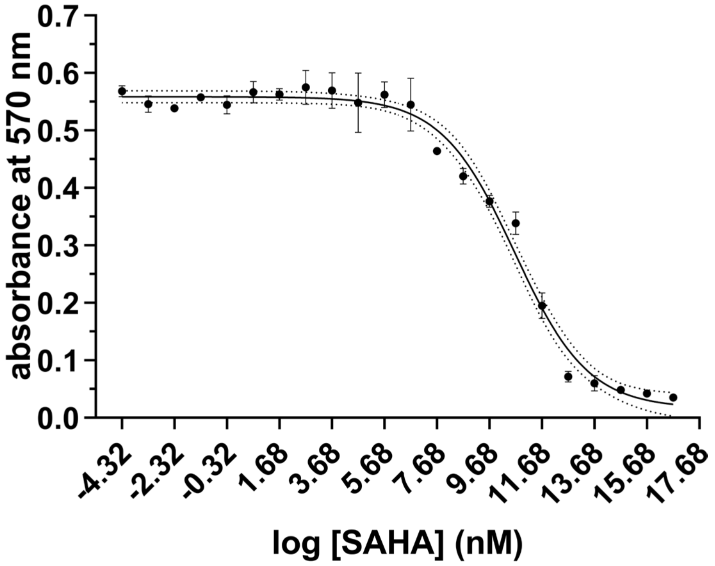
